# The stress-induced lincRNA *JUNI* is a critical factor for cancer cell survival whose interactome is a prognostic signature in clear cell renal cell carcinoma

**DOI:** 10.1101/2023.10.23.563579

**Authors:** Vikash Kumar, Xavier Sabaté-Cadenas, Isha Soni, Esther Stern, Carine Vias, Doron Ginsberg, Carlos Romá-Mateo, Rafael Pulido, Martin Dodel, Faraz K. Mardakheh, Iddo Z. Ben-Dov, Alena Shkumatava, Eitan Shaulian

## Abstract

Cancer cells rely on adaptive mechanisms to survive the multiple stressors they encounter, including replication stress, toxic metabolic products and exposure to genotoxic drugs. Understanding the factors involved in these stress responses is crucial for developing effective treatments. Here, we describe a previously unstudied long non-coding RNA (lncRNA), *JUNI* (*JUN-DT*, *LINC01135*), which is regulated by MAPK and responsive to stress. *JUNI* positively regulates the expression of its neighboring gene *JUN*, a key transducer of signals that regulate multiple transcriptional outputs. Our findings reveal that silencing *JUNI* sensitizes cancer cells to chemotherapeutic drugs or UV radiation, and that its prolonged silencing leads to cell death regardless of stress exposure, highlighting the pro-survival importance of *JUNI.* We identified 57 proteins that interact with *JUNI* and found that the activity of one of them, the MAPK phosphatase and inhibitor DUSP14, is inhibited by *JUNI*. This effect results in c-Jun induction following exposure of cancer cells to UV radiation and promotes cellular survival. Although *JUNI* regulates c-Jun and its downstream targets, the pro-survival effects in cells not exposed to stress are only partially dependent on c-Jun regulation.

*JUNI* expression levels significantly correlate with patients survival across 11 different types of cancer. Interestingly, the correlation of DUSP14 expression levels with patients survival in nine of these tumors is coherently inverse, indicating contradicting effects that are relevant not only for c-Jun induction and cellular survival but also in human cancer. Notably, we observed particularly significant antagonistic correlations in clear cell renal cell carcinoma (ccRCC) (p=5.7E-05 for *JUNI* and p=2.9E- 05 for Dusp14). In fact, the expression levels of 76% of *JUNI*-interacting proteins predict the prognosis of ccRCC patients significantly. Furthermore, a combined hazard ratio calculation demonstrates that this gene combination serves as a highly specific prognostic signature for ccRCC. Overall, our findings reveal a new important factor in stress signaling and cellular survival that is involved in ccRCC.

## Introduction

Long noncoding RNAs (lncRNAs) have emerged as important players in the neoplastic process due to changes in their expression and activity, as supported by functional experimental evidence (Bhan *et al*, 2017; Chi *et al*, 2019; Spizzo *et al*, 2012; Tan *et al*, 2021). LncRNAs have been shown to affect various cancer-related processes and can impact every stage of cancer development, including initiation, progression, and metastasis (Gutschner & Diederichs, 2012; McCabe & Rasmussen, 2021; Tan *et al*., 2021). Specifically, the involvement of lncRNAs in stress responses, particularly the DNA damage response (DDR), allows cancer cells to regulate and overcome DNA damage and associated cellular responses, which can otherwise lead to cell death. As such, lncRNAs play a crucial role in promoting cancer cell survival and chemoresistance to therapy (Chen *et al*, 2017; Connerty *et al*, 2020; Su *et al*, 2018; Tehrani *et al*, 2018).

Rapid DNA damage responses (DDRs) are critical for the maintenance of organismal wellbeing and are often disrupted during cancer development (Jackson & Bartek, 2009). Tumor suppressors such as ATM, p53, and BRCA proteins are potent participants in the sensing, transduction, and repair of DNA damage (Jackson & Bartek, 2009). Conversely, some oncoproteins, including the AP-1 member c-Jun, are also responsive to stress signaling and are particularly sensitive to environmental DNA-damaging agents such as ultraviolet light (UV) (Devary *et al*, 1991). UV-induced phosphorylation of c-Jun N-terminal kinase (JNK) leads to JNK-dependent c-Jun phosphorylation, protein stabilization, and elevated transcription due to autoregulation of its expression (Davis, 2000; Fuchs *et al*, 1998; Karin & Gallagher, 2005; Stein *et al*, 1992). The wide responsiveness of c-Jun to environmental stimuli and its consequential multiple roles in cellular homeostasis make it essential for cellular survival (Shaulian & Karin, 2002), development (Hilberg *et al*, 1993) and tumorigenicity (Eferl *et al*, 2003). Moreover, the ability of c-Jun to antagonize proapoptotic signaling underlies its capacity to endow cancer cells with drug resistance. For example, its activation post-treatment of melanomas with BRAF inhibitors is a primary driver for phenotype switching, rendering the cells more mesenchymal and associated with elevated drug resistance (Fallahi-Sichani *et al*, 2015; Ramsdale *et al*, 2015; Titz *et al*, 2016).

In 2020, around 430,000 new cases of kidney cancer and 180,000 deaths were reported worldwide, with clear cell renal cell carcinoma (ccRCC) accounting for approximately 75% of cases (Hsieh *et al*, 2017; Sung *et al*, 2021). The first comprehensive characterization of ccRCC re-verified the loss of chromosome 3p encompassing VHL, PBRM1, SETD2 and BAP1. It also identified 19 significantly mutated genes, epigenetic alterations including methylation of specific promoters and widespread DNA hypomethylation associated with mutations in SETD2, recurrent mutations in the PI3K/AKT pathway and aberrations in the chromatin remodeling complex SWI/SNF (Cancer Genome Atlas Research, 2013). The array of genes, which are mutated, inactivated, or hyper-activated in renal tumorigenesis, are reported to be involved in the regulation of various metabolic events, such as glycolysis, TCA cycle, metabolism of glutamine, ATP production and modulation of pathways important for hypoxic condition and redox balance (Rathmell *et al*, 2018; Weiss, 2018; Wettersten, 2020). The essential genes involved in regulating metabolic reprogramming in renal cancer are VHL, PTEN, Akt, mTOR, TSC1/2 and Myc (Schmidt & Linehan, 2016). Inactivation of VHL leads to the stabilization of two VHL E3 ubiquitin ligase complex targets, HIF1α and HIF2α (Shen & Kaelin, 2013). Under hypoxic state of cancer cells, HIF1α and HIF2α up-regulate the transcription of several hypoxia-responsive genes involved in tumor growth, angiogenesis, and metastasis, as well as genes associated with glucose transport and metabolism (Kaelin, 2002, 2004) In addition, altered fatty acid metabolism is observed in ccRCC. In fact, accumulation of “lipid droplets” are considered to be a hallmark of clear-cell renal cell carcinoma (ccRCC) as these lipid droplets in the cytoplasm give rise to the typical clear cell phenotype (Wettersten *et al*, 2015). In the recent years the roles of several lncRNAs serving as oncogenes or tumors suppressors in tumorigenesis of renal cell carcinoma is revealed (Shen *et al*, 2021).

In this study, we identified *JUNI* as a stress-regulated lncRNA that controls c-Jun expression. We show its requirement for the survival of cancer cells and suggest a possible role in ccRCC.

## Materials and methods

### Cell culture

CHL-1, HeLa, MDA-MB 231, HACAT and HMCB cells were procured from ATCC. The first 4 were cultured in DMEM with 10% FBS, whereas HMCB cells were cultured in EMEM. All cells were grown with 10% FBS and cultured at 37°C with 5% CO_2_. Cells were routinely monitored for mycoplasma contamination.

### Transfection, treatments, and inhibitors

Plasmids were transfected into 40-50% confluent cells using PloyJet reagent (SignaGen Lab MD, USA). siRNAs were transfected with the siRNA transfection reagent INTERFERin® (Polyplus, Illkirch – France). The siRNA concentrations used in this study ranged between 5 and 20 nm. SP600125 (JNK) or SB203580 (p38) inhibitors (10 µM) were added one hour prior to UV irradiation. Doxorubicin, etoposide and cisplatin were used at doses of 0.5 to 5 µM. XTT viability assays (Abcam, Cambridge UK) were used according to the manufacturer’s instructions.

### RNA Extraction and Real-Time PCR

RNA was isolated manually using TRI Reagent® (MRC, OH, USA). cDNA was prepared from total RNA (1 µg) using Quantabio, and a qScript^TM^ cDNA synthesis kit and PerfeCTa SYBR Green SuperMix were used for qPCR according to the manufacturer’s instructions (Quantabio, MA, USA). To prepare DNA-free RNA samples, RNA was treated with PerfeCTa® DNase I according to the manufacturer’s instructions (Quantabio, MA, USA). A StepOne Plus Real-Time PCR apparatus with StepOne Software v2.3 was used for analysis (Applied Biosystems, MA USA). Each experiment was performed in multiple technical duplications (n>4) with at least 3 biological repeats.

### Calculation of JUNI copy number

RNA was extracted from 1.5X10^5^ HMCB and MDA-MB-231 cells, cDNA was prepared and qPCR performed with primers aimed at the first exon of *JUNI* as described above. To calculated the total amount of *JUNI*/well CT values of *JUNI* RNA obtained from both cell lines were compared to calibration curve of CT values (y) of known amounts of *JUNI* expressing plasmid (X). Number of copies/well (NCW) was calculated using the following equation: NCW=(ng×[6.022×10^23^])/(length ×[1×10^9^]×650). Number of copies/cell (NCC) was calculated by the following formula: NCC=NCW*dilution factor from total RNA to cDNA in a single well/number of cells from which the RNA was extracted. Taken into account 20-30% loss of RNA in extraction and RTqPCR process the number of copies presented is the minimal copy number per cell.

### Antibodies, immunobloting and proteins quantigication

The primary antibodies used were c-Jun (CST #60A8 1:1000), Ser-63-p-JUN (CST #9261 1:500), actin (CST #3700S 1:5000), GAPDH (CST #97166 1:5000), P-JNK (CST #9251 1:500), total JNK (#9252, #9258 1:1000), cleaved Caspase-3 Asp175 (5A1E #96641:1000) and anti-FLAG M2 antibody (Sigma, #F1804). HRP-conjugated secondary antibody was used at a dilution of 1:5000 (Jackson Laboratories, USA). Immunobloting was performed using standard protocol. Proteins were detected and quantitated using Bio-Rad’s chemiDoc imagers. Representative images are presented. Proteins quantities normalized to loading controls are presented below each relevant blot. In all cases N≥3.

### Colony formation, soft agar assay and spheroid generation

A total of 1000-2000 cells were seeded in 3.5 cm dishes after different transfections. The cells were fixed 12-14 days later with a methanol:acetic acid (3:1) ratio, and colonies were stained with methylene blue (1% methylene blue in 0.1 M borate buffer, pH 8.5) and counted. Low amounts (50-100 ng/well in 12 well dish) of *JUNI* expression vector were transfected 24h and 120h post siRNA transfection in rescue experiments. Transfection efficiency of DNA was calculated by GFP staining, and the number of colonies was normalized to transfection efficiency. For soft agar assays, 10k-15k cells were seeded on 1% agar/complete media mixture-coated 3.5 cm plates and covered with 0.3% agar/complete media mixture, and the cells were incubated for 15-20 days. For spheroid generation, 2-3k cells were seeded using hanging drop method, 5-7 days later spheroids were transferred to ultra low-attachment U-bottom 96 well plate and treated with 1-2 µM of doxoruicin for 5-8 days. Images were acquired at 10X magnification, spheroids were trypsanized and total cells number was counted using Trypan blue dye.

## Microscopy

Images were captured at 20X using an inverted Nikon Eclipse T2S microscope. Scale bars are presented.

### Plasmids and cloning

Plasmids expressing *JUNB*, *JUND* and mutant *JUN* that cannot bind DNA (272/273E) were previously described (Yogev *et al*, 2008). DUSP14-HIS was a kind gift from. M. Saleh (Douglas & Saleh, 2019) (Douglas & Saleh, 2019) pRK5 HA-DUSP14 plasmid (HA tag N-terminal) was obtained by PCR amplification from pOTB7 DUSP14 plasmid (HGMP MRC geneservice, IMAGE cDNA clone 2819474) and subcloning into pRK5 plasmid containing HA sequence. pRK5 HA-DUSP14 C111S was obtained by PCR oligonucleotide site-directed mutagenesis. To generate the *JUNI*-MS2×10 construct used in incPRINT, a 938 bp fragment of the *JUNI* mature RNA was cloned into 10xMS2 vector (Graindorge *et al*, 2019) using a gBlocks Gene Fragment containing Cla1 and Hind3 restriction sites (IDT, NJ, USA). Exon1 of *JUNI* was cloned into pEFA1 CMV puro/GFP vector using a gBlocks gene fragment containing EcoR1 and BAMH1 restriction enzymes. The genomic element that contains the promoter of *JUN* flanked by 153 bases of the first exon of *JUNI* on one side and 750 bp of the 5’ UTR of *JUN* on the other side as well as its mutant versions were previously described (Angel *et al*, 1988).

### Crosslinking and Immunoprecipitation (CLIP)

Ten million HeLa cells were transiently transfected with mammalian expression vectors expressing His-DUSP14 or GFP as a control using Lipofectamine 2000 (Thermo Fisher) according to the manufacturer’s instructions. The next day, the cells were washed with PBS and irradiated once on ice with 150 mJ/cm^2^ UV light (254 nm) in ice-cold PBS using a Hoefer Scientific UV Crosslinker. Cells were then pelleted and lysed in 1 ml of lysis buffer (50 mM Tris-HCl pH 7.4, 100 mM NaCl, 1% Igepal CA-630, 0.1% SDS, 0.5% sodium deoxycholate, supplemented with protease inhibitors), sonicated, cleared, and adjusted to a protein concentration of 1 mg/ml. RNA was then partially digested with 0.2 U/ml RNase I for 3 mins before immunoprecipitation with 10 μL of anti-His-tag antibody (Cell Signaling) pre-conjugated to 50 μL protein G Dynabeads (Thermo Fisher) for 1 hour. The beads were then washed 5 times with lysis buffer before the bound RNAs were eluted off by Proteinase K (Thermo Fisher) digestion for 1 hour at 65°C. The RNA was then extracted using TRIzol (Thermo Fisher) and subjected to qPCR analysis with qPCR primers against *JUNI*, *MALAT1*, and *PVT1*.

### IncPRINT

The incPRINT protocol and analysis were previously described (Graindorge *et al*., 2019). In this study, the bait was a 938 bp fragment of the *JUNI* mature RNA cloned into the 10xMS2 vector. The experiment was performed twice. The threshold for significant interaction was set at a relative luciferase activity (protein versus background) 2.2 or higher.

### Statistical analysis of JUNI interacting genes in ccRCC

Median survival durations associated with high and low tumor expression of each of the *JUNI*- interactors in patients with ccRCC (KIRC) or bladder cancer (BLCA) and associated logrank test p-values were extracted from Pan-cancer RNA-seq Kaplan-Meier plotter (Nagy *et al*, 2021). We then calculated the log_2_ values of the median survival ratios (MSR), defined as median survival in patients with high expression divided by the median survival in patients with low expression.

The mortality hazards ratios (HRs) associated with 1 standard deviation (SD) increase in tumor expression of each of the *JUNI*-interactor genes in patients of all Cancer Genome Atlas (TCGA) cohorts were derived by downloading expression and survival data from the Broad GDAC Firehose repository, and processing as previously described. [https://www.biostars.org/p/153013/ (Parnell *et al*, 2011)]. Next, a collective expression score was calculated by adding the expression z-values of *JUNI* interactors associated with worse prognosis and subtracting the expression z-values of interactors associated with improved prognosis, based on mortality prediction in the KIRC cohort. We used this score as an independent mortality predictor in Cox proportional hazards models in each TCGA cohort.

### Primers and siRNA

### PCR primers

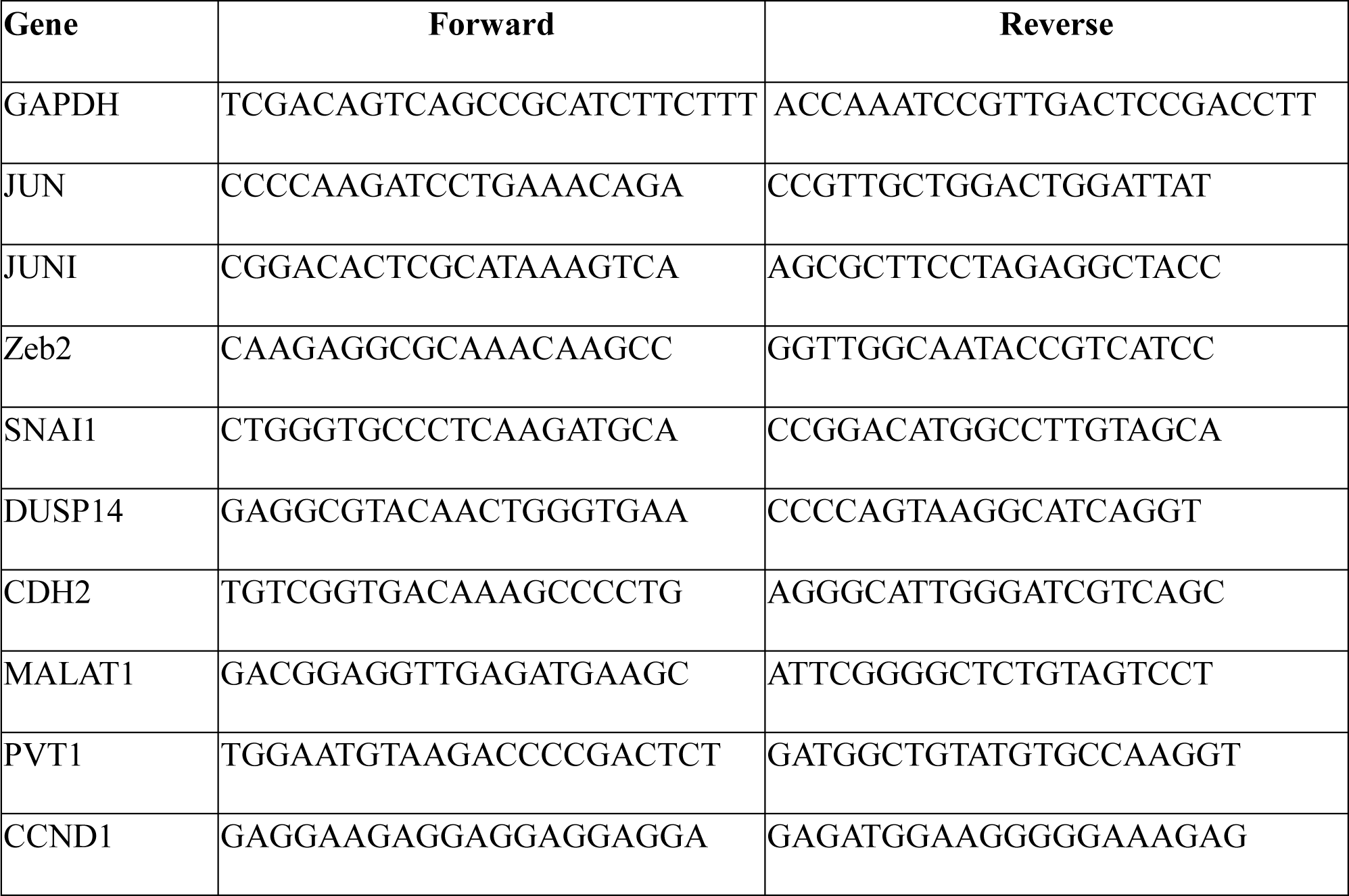

### siRNA oligonucleotides

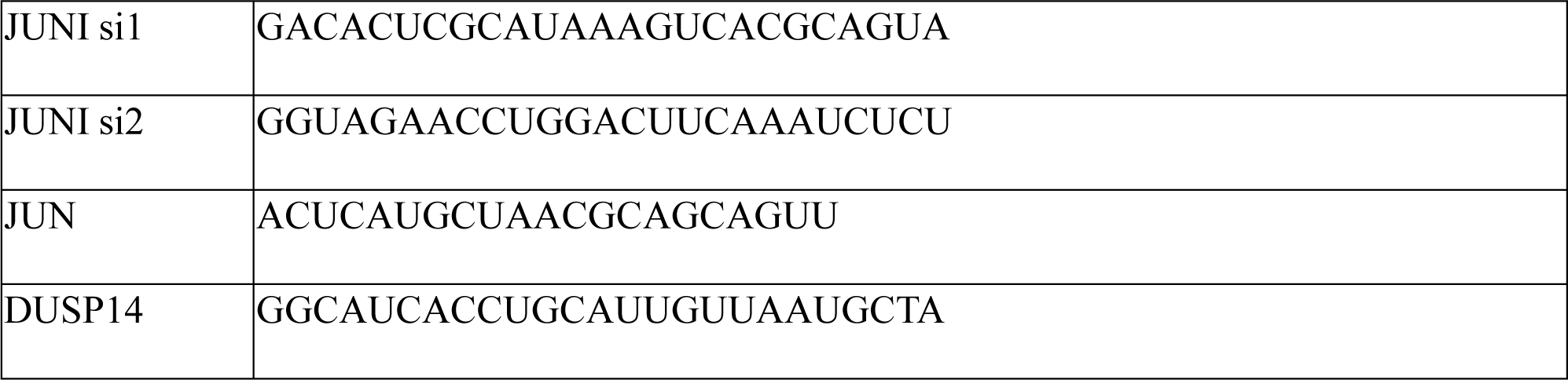

## Results

### *JUNI* is a stress-regulated lincRNA

To identify genes regulated by c-Jun, we analyzed ChIP-seq data from ENCODE for genes whose transcription start sites (TSSs) are located in the vicinity of c-Jun binding sites. One of the genes whose TSS is located only 1100 bp away from the *JUN* TSS is the long intergenic noncoding RNA *JUN-DT* or *LINC01135*, referred to hereafter as *JUNI* (for *JUN* inducer). According to ENCODE data, *JUNI* contains five main exons and has multiple isoforms. Twenty-seven different transcript isoforms were described according to LNCipedia ranging from 213 to 6213 bases (Volders *et al*, 2019). Importantly, ENCODE predicts that the first exon is shared by all, therefore, all primers to analyze JUNI’s expression as well as siRNAs to silence it, were targeted for this exon. *JUNI* is evolutionarily conserved within primates but not in rodents (Fig 1A). The fact that c-Jun is known to bind to its own promoter and to autoregulate its expression (Angel *et al*., 1988; Stein *et al*., 1992) supported the ChIP-seq results but did not confirm any functionality for *JUNI*. Therefore, we analyzed its expression to exclude promoter leakiness. UV-driven activation of c-Jun N-terminal kinase (JNK) results in JNK-dependent c-Jun phosphorylation, transcript elevation and protein stabilization (Davis, 2000; Fuchs *et al*., 1998; Karin & Gallagher, 2005). Hypothesizing that the expression of *JUNI* is regulated by c-Jun and given that UV radiation is a major inducer of c-Jun expression, we irradiated four different cell types with UV radiation and examined *JUNI* expression. In this study, we used a set of cell lines, including two melanoma cell lines (where UV is a major carcinogen), HMCB and CHL1, HeLa (cervical carcinoma cell line in which *JUN* expression was intensively studied) and MDA-MB-231 (breast cancer-TNBC; a cancer type in which *JUN* plays a significant role) (Vleugel *et al*, 2006). Irradiation of these cell lines with 20-30 J/m^2^ UVC resulted in three to five-fold induction of *JUNI* expression (Fig 1 B). Similar to *JUN*, the induction was dose dependent (Fig 1C), and the rapid response to stress (Fig 1D) as well as to serum stimulation of starved cells, identified by others (Bunch *et al*, 2019), qualifies it as an “immediate early” lncRNA. To determine how common is the correlation between *JUNI* and *JUN* induction following exposure to drug-induced stress, HeLa cells were exposed to chemotherapeutic drugs that induce *JUN* expression, and *JUNI* levels were monitored. Every DNA damaging treatment that induced *JUN* also induced *JUNI*, although to a lower extent (Fig 1E). Fractionation experiments demonstrated that *JUNI* resides mainly in the nucleus (Fig. 1F) and quantitation of *JUNI*’s copy number in untreated HMCB and MBA-MD-231 cells revealed the presence of of about 8 copies per cell. HeLa cells express lower amounts of 1-2 copies in untreated cells. Overall, these results suggested that *JUNI* is a stress-induced gene whose expression pattern resembles that of *JUN*, therefore, we investigated the potential existence of regulatory effects between the two genes, especially post exposure of cells to stress.

**Figure 1.**
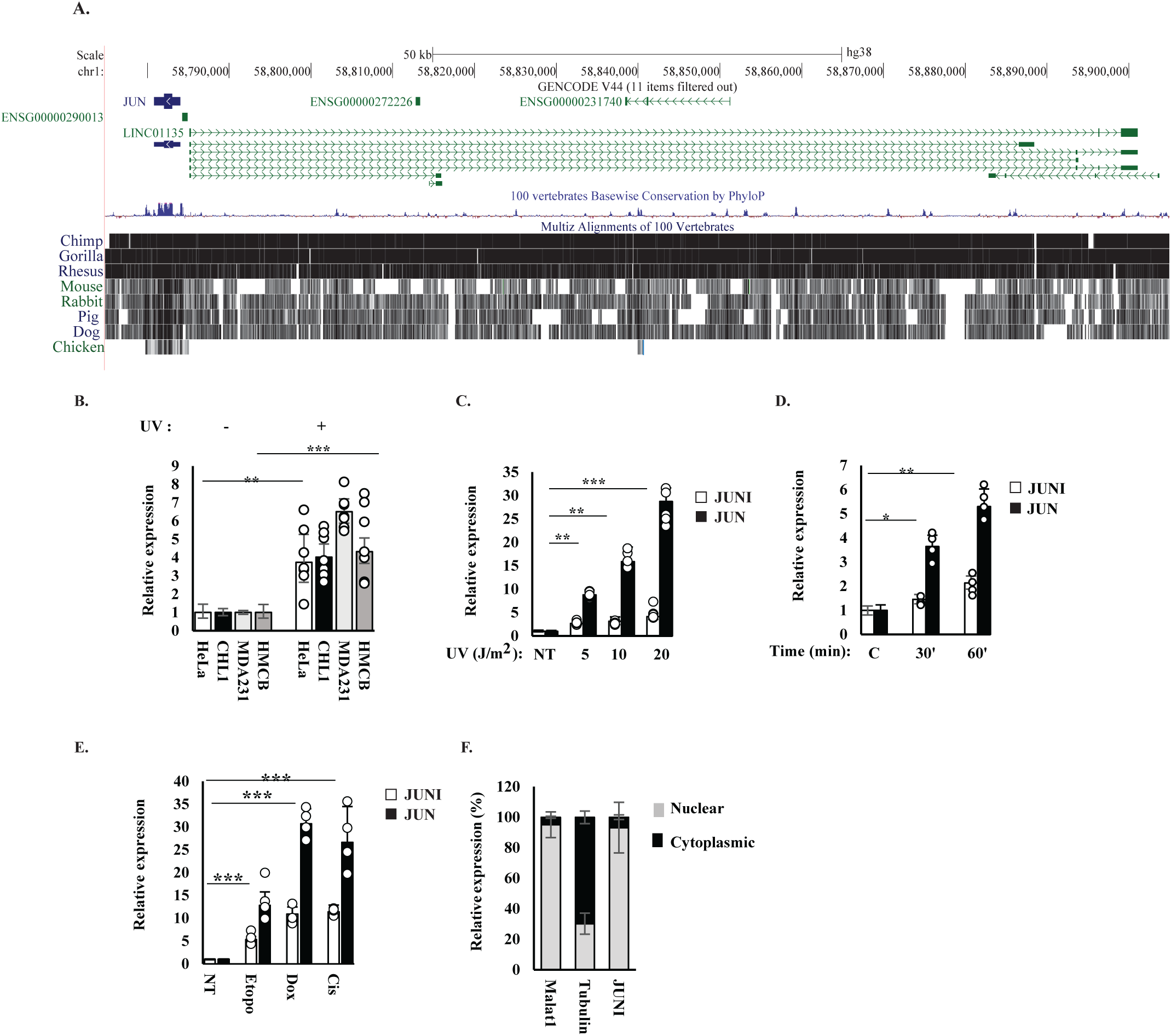
*JUNI* is a nuclear lncRNA upregulated by stress. A. A schema of *JUNI* structure and evolutionary conservation. Green lines represent *LINC01135* isoforms (*JUNI*), black lines alignment with the indicated species. Data were obtained from the UCSC Genome Browser on Human (GRCh38/hg38). B. Expression of *JUNI* in HeLa, CHL1, MDA-MB-231 and HMCB cells before and 4 h after exposure to 20 or 30 J/m^2^ UV (HeLa and all other cell lines, respectively). GAPDH was used for normalization in this experiment and in all the following qPCR experiments. C. HeLa cells were irradiated with the indicated doses of UV and harvested 4 h after irradiation. *JUNI* and *JUN* RNA levels (white and black bars, respectively) were determined by RT-qPCR. D. HeLa cells were irradiated with 20 J/m^2^ UV and harvested 30 or 60 min after irradiation. *JUNI* and *JUN* RNA levels (white and black bars, respectively) were determined by RT-qPCR. E. HeLa cells were untreated (NT) or exposed to 5 μM of etoposide, doxorubicin or cisplatin. *JUN* and *JUNI* RNA levels (black and white bars, respectively) were determined 4 h after treatment using RT-qPCR. F. HeLa cells were fractionated into nuclear and cytoplasmic fractions, and the abundance of *JUNI* was examined in each fraction using RT-qPCR. The gray portion of the bar represents the nuclear fraction, and black represents the cytoplasmic fraction. Tubulin and MALAT1 RNAs were used as markers for cytoplasmic and nuclear fractions, respectively. Significance is marked in this study according to the following key: *, p<0.05; **, p<0.001, ***, p<1×10^-5^. N in all RT-qPCR experiments was >3.

### *JUNI* expression is controlled by MAPKs but not by c-Jun

To define the pathways required for *JUNI* upregulation after stress exposure, we inhibited two UV-activated kinases that are known to phosphorylate/induce c-Jun, JNK and p38 using specific inhibitors SP600125 (JNK) or SB203580 (p38). Both inhibitors abolished *JUNI* induction by UV, similar to c-Jun expression (Fig 2 A, B), suggesting that the expression of *JUNI* may be c-Jun-dependent. As the *JUN* promoter is the major regulatory element proximate to the first exon of *JUNI,* we tested whether it co-regulates *JUNI* expression. We transfected MDA-MB-231 and HeLa cells with a genomic element that contains the promoter of *JUN* flanked by 153 bases of the first exon of *JUNI* on one side and 750 bp of the 5’ UTR of *JUN* on the other side (Angel *et al*., 1988). Examination of RNA extracted after DNase treatment demonstrated eight-and 34-fold higher expression of *JUNI* relative to cells transfected with an empty vector in the different cell lines (Fig 2C and D). Thus, *JUN*’s promoter is bidirectional, driving the expression of *JUN* on one side and *JUNI* on the other, as previously described for other lncRNAs (Kopp & Mendell, 2018). Two AP-1 binding sites in the *JUN* promoter mediate autoregulation of *JUN* expression. Surprisingly, although mutation in these sites reduced *JUN* expression relative to the promoter expressing wt AP-1 binding sites, *JUNI* expression levels did not change, suggesting that *JUNI* is not regulated by c-Jun (Fig 2D). To directly explore the dependence of *JUNI* expression on c-Jun, specific siRNA for *JUN* was transfected into HeLa cells, the cells were UV-treated or not, and the levels of c-Jun and *JUNI* were determined (Fig 2E). Despite three-fold suppression of c-Jun protein expression*, JUNI* levels did not decline (Fig 2E). Reciprocal experiments supported the independence of *JUNI* from c-Jun expression. HeLa cells transfected with the *JUN* expression vector and expressing very high levels of *JUN* mRNA did not present any difference in *JUNI* expression (Fig 2F), and unlike the induction of *CCND1,* a known c-Jun target gene (Wisdom *et al*, 1999), *JUNI* levels were not changed in the *JUN*- transfected cells expressing high levels of c-Jun protein (Fig 2G), thus proving that *JUNI* expression is c-Jun independent.

**Figure 2.**
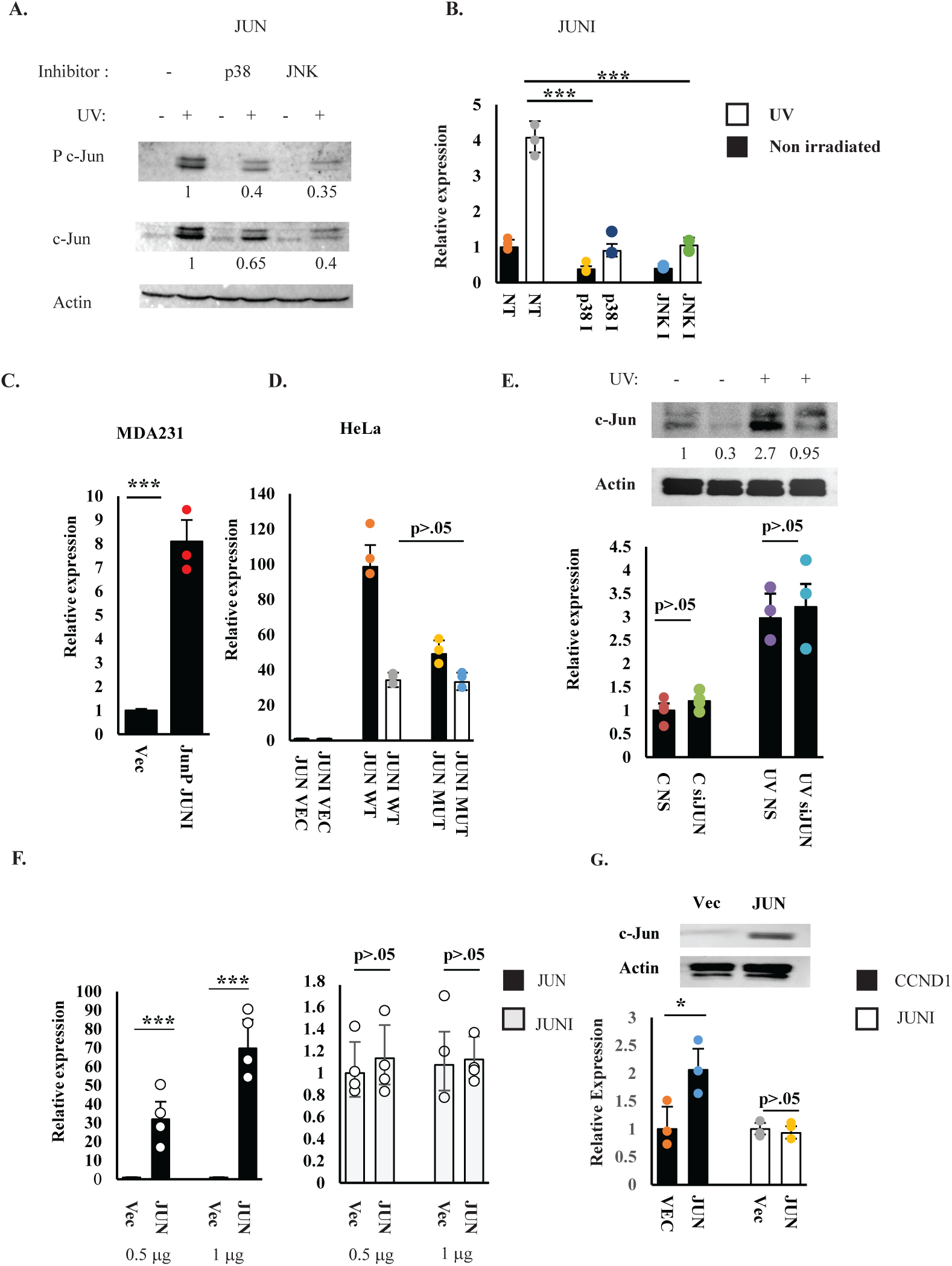
*JUNI* induction by UV is MAPK-dependent and c-Jun independent. HeLa cells were treated with 10 mM SP600125 (JNK inhibitor) or SB203580 (p38 inhibitor) 1 h prior to exposure to 20 J/m^2^ UV and harvested 2 h after exposure. Phosphorylated (Ser 63) and total c-Jun protein levels were determined using specific antibodies, (A) and *JUNI* RNA levels were determined using RT-qPCR (B). Numbers under each panel of western blots indicate normalized relative quantities of the the relevant protein. White bars indicate RNA from UV-exposed cells, and black bars indicate RNA from nonexposed cells. Actin was used as a loading control. C. MDA-MB-231 HeLa cells were transfected with a vector containing the genomic sequences of the *JUN* promoter and the adjacent first exon of *JUNI* or with an empty vector. Forty-eight hours later, *JUNI* expression was monitored using RT-qPCR after DNase treatment. D. HeLa cells were transfected with the same vector or with a vector containing mutated AP-1 binding sites in the *JUN* promoter. Levels of *JUN* (black bars) or *JUNI* (white bars) were determined using RT-qPCR. E. HeLa cells were transfected with nonspecific (NS) or *JUN*-specific siRNAs (10 nM), irradiated with 20 J/m 48 h post transfection or not, and harvested 3.5 h later. c-Jun protein (upper panel) and *JUNI* levels (lower panel) were measured using immunoblotting and RT-qPCR. F. HeLa cells were transfected with the indicated amounts (micrograms) of empty vector or c-Jun expression vector, irradiated 36 h later with 20 J/m^2^ UV and harvested 6 h after irradiation. The levels of *JUN* RNA (black bars) or *JUNI* (gray bars) were measured using RT-qPCR. G. HeLa cells were transfected with 500 ng of empty vector or c-Jun expression vector. Cells were harvested 48 h later, and c-Jun protein levels (upper panel) or *CCND1* (black bars) and *JUNI* (white bars) RNA levels, were measured using immunoblotting and RT-qPCR, respectively. Al westerns in this experiment and all the following figures N≥3.

### *JUNI* regulates c-Jun expression and its downstream targets and is relevant for human cancer

Given that many lncRNAs modulate the expression of nearby genes in a *cis* (Gil & Ulitsky, 2020), we examined the effects of *JUNI* on *JUN* expression. To this end, we attempted to knock out the first exon of *JUNI* using CRISPR gene editing targeting *JUNI* with two sgRNAs flanking its first exon. However, exon 1-deleted clones could not be generated, suggesting an essential role for this gene. As the first exon is common to all isoforms, we designed two siRNAs targeting it and used them to transiently silence *JUNI* expression. Both siRNAs silenced *JUNI* and consequently *JUN* mRNA expression to a lesser degree (Fig. 3A). It is worth mentioning that cell line-specific, consistent variations in the silencing capacities of the two siRNAs were observed. Importantly, all the phenotypic effects described in this study corresponded to the silencing efficiencies of each siRNA. HMCB and MDA-MB-231 cells express high levels of c-Jun protein, which can be detected even without stress exposure. A mild reduction in c-Jun protein levels after *JUNI* silencing was detected in these cells (Fig 3B). In contrast, a clear effect of silencing was observed in all cell lines after exposure to UV (Fig 3C), suggesting that *JUNI* has special importance for c-Jun expression after exposure of cells to stress. Indeed, exogenous expression of the first exon of *JUNI* either transiently (Fig 3 D) or stably (Fig 3E) was sufficient to elevate c-Jun levels in cells exposed to UV radiation. Combined, our data suggest that *JUNI* can regulate *JUN* expression in *trans* and is particularly important after stress exposure. Significant correlations between *JUNI* and *JUN* expression also occur *in vivo*. Examination of expression data of 32 types of tumors in the Pan-Cancer Co-Expression Analysis for the RNA-RNA interactions (Li *et al*, 2014) revealed a 94% positive correlation between *JUNI* and *JUN* levels in all cancer types examined. Sixty-two percent of the positively correlated cases were statistically significant (Table S1).

**Figure 3.**
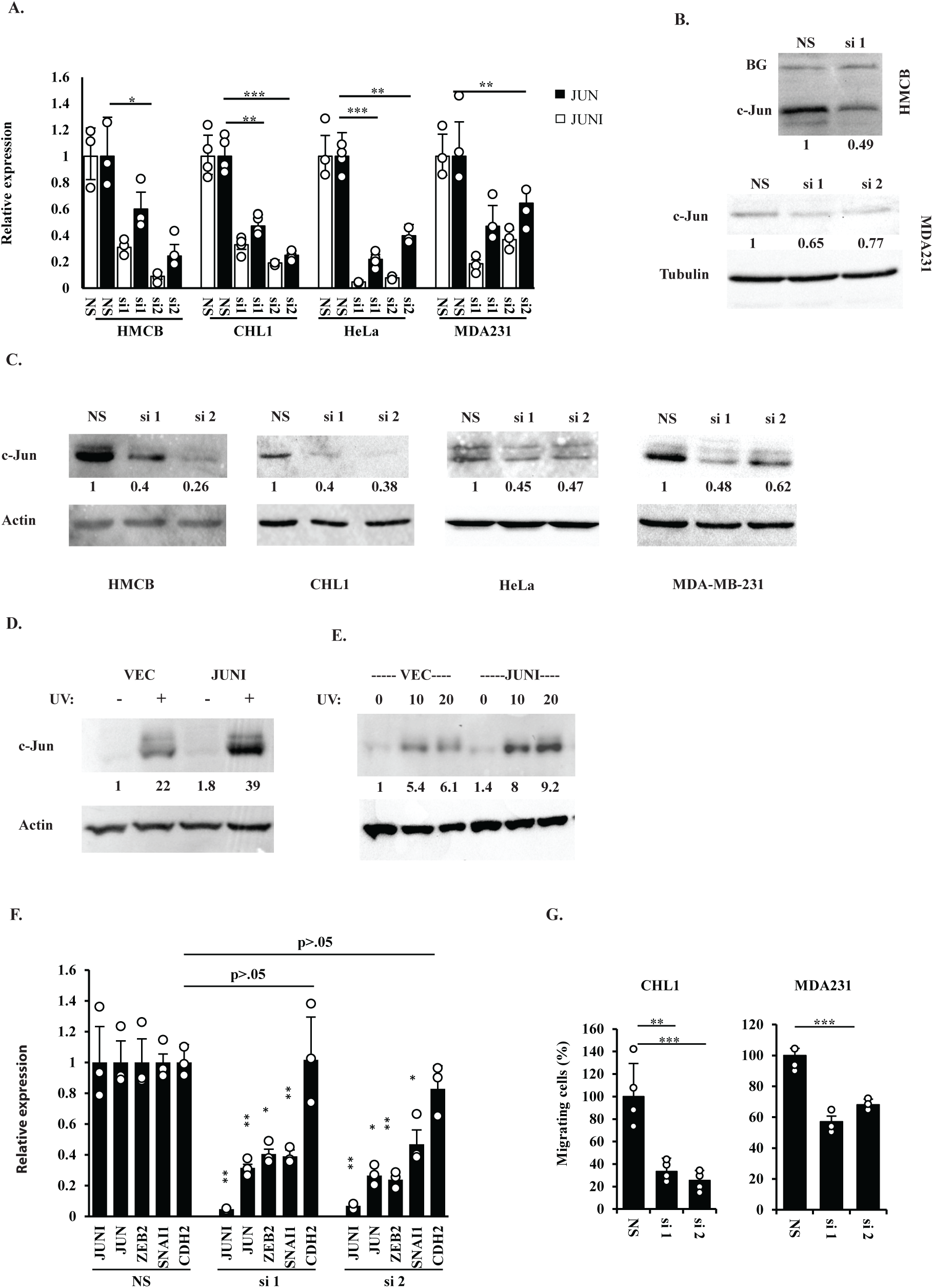
*JUNI* regulates the expression of *JUN*, its target genes and cellular motility. A. The indicated cells were transfected with nonspecific siRNA (NS) or two siRNAs against JUNI (si1: 20 nM and si2: 5 nM). The cells were harvested 24 h later, and RT-qPCR was performed to determine *JUNI* (white bars) and *JUN* (black bars) levels B. HMCB cells (upper part) and MDA-MB-231 cells (lower part) were transfected with NS-or *JUN*I-specific siRNAs and harvested 48 h later. c-Jun protein levels were determined by immunoblotting using a specific antibody. Equal background bands (BGs) or actin were used as loading controls. C. The indicated cells were transfected with NS-or *JUNI*-specific siRNAs and exposed to UV radiation. c-Jun protein levels were examined by immunoblotting 4-6 h after exposure using a specific antibody. Actin was used as a loading control. HeLa cells were transiently (D) or stably transfected (E) with the first exon of *JUNI* or with an empty vector. Cells were irradiated 36 h post transfection with 20J/m^2^ (D) or with the indicated UV doses, and the levels of c-Jun protein were measured 4 h later as described above. F. CHL1 cells were transfected with the indicated siRNAs. Expression of the indicated genes was determined 24 h later using RT-qPCR and specific primers. G. The percentage of siRNA-transfected cells that crossed the 8-micron Millicell hanging insert membrane within 16 hours post plating. A total of 500000 cells were seeded 24 h after transfection with the indicated siRNA. The migration of NS siRNA-transfected cells was considered 100%. The mean plus SD is presented. N: CHL1, 4; MDA-MB-231, 3.

Next, we examined whether *JUNI* has a regulatory impact on targets downstream of c-Jun. As c-Jun can modulate melanoma cells plasticity (Ramsdale *et al*., 2015), we examined the expression of a set of known c-Jun-regulated genes in CHL1 cells 24 h after silencing *JUNI*. Interestingly, c-Jun target genes known to be involved in EMT, such as ZEB2 (Graindorge *et al*., 2019; Zhao *et al*, 2014) and SNAI1 (Thakur *et al*, 2014), were downregulated by *JUNI* silencing, whereas CDH2, which is not a known target, was not (Fig 3 F). The effect on these EMT-relevant *JUN* targets was also associated with reduced motility, as reflected by the capacity of cells to migrate in transwell migration assays (Fig. 3G and S1).

To investigate the potential relevance of *JUNI* in human cancer, given its regulatory impact on the neighboring *JUN* gene and its influence on motility, we analyzed the correlation between *JUNI* expression levels and patients survival in 21 different types of cancer with available RNA-seq data from the Pan-cancer Project (Nagy *et al*., 2021). Our analysis revealed significant correlations between *JUNI* expression and patient survival in 11 of the cancer types examined, either negatively or positively depending on the specific cancer type (Table 1).

**Table 1:**
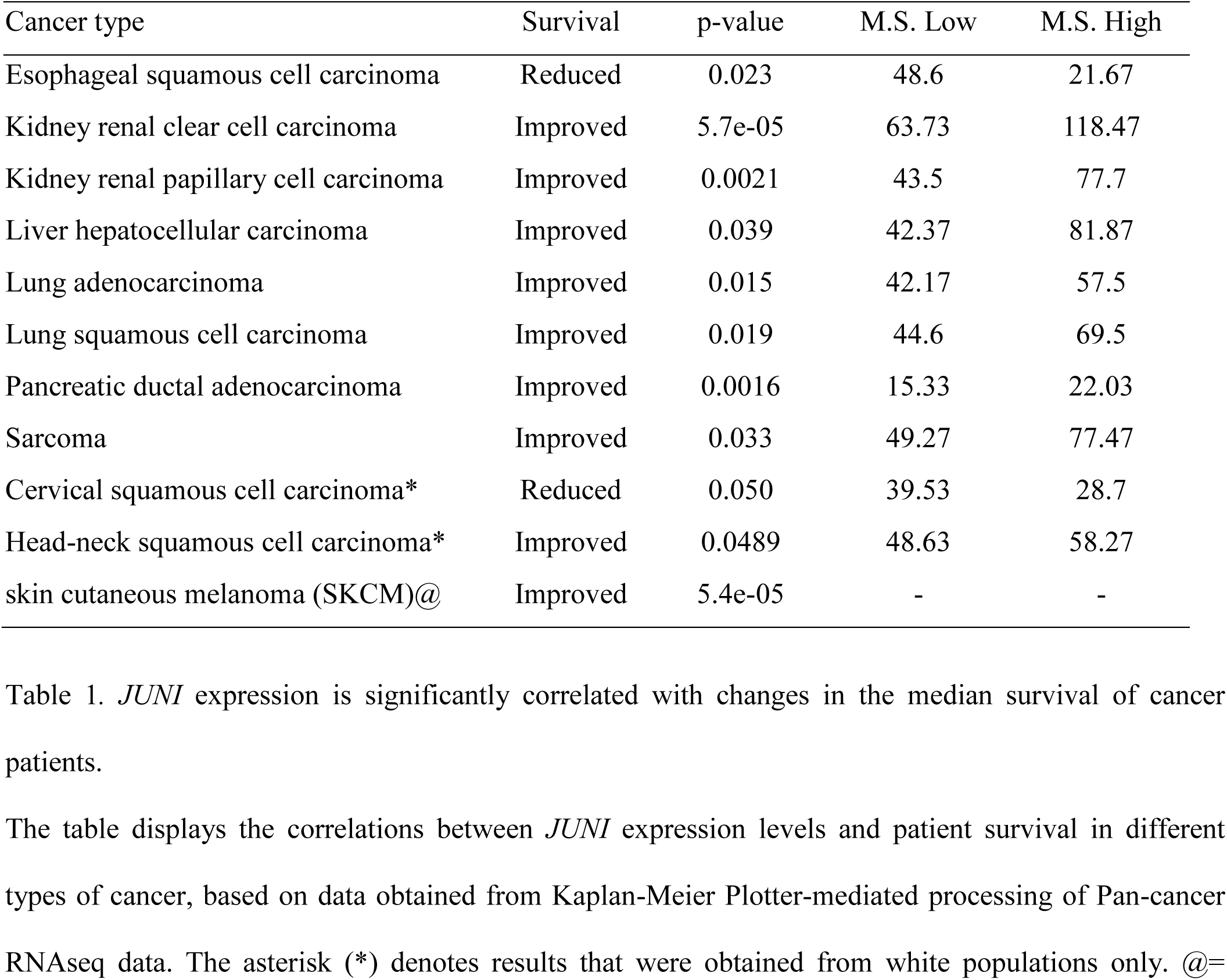
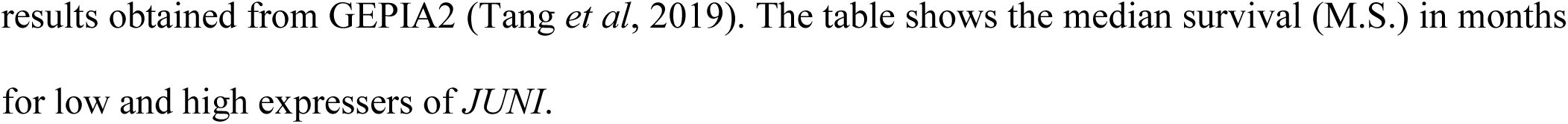
Association of JUNI expression with median survival in cancer patients.

JUNI* expression is significantly correlated with changes in the median survival of cancer patients.
| Cancer type | Survival | p-value | M.S. Low | M.S. High |
| --- | --- | --- | --- | --- |
| Esophageal squamous cell carcinoma | Reduced | 0.023 | 48.6 | 21.67 |
| Kidney renal clear cell carcinoma | Improved | 5.7e-05 | 63.73 | 118.47 |
| Kidney renal papillary cell carcinoma | Improved | 0.0021 | 43.5 | 77.7 |
| Liver hepatocellular carcinoma | Improved | 0.039 | 42.37 | 81.87 |
| Lung adenocarcinoma | Improved | 0.015 | 42.17 | 57.5 |
| Lung squamous cell carcinoma | Improved | 0.019 | 44.6 | 69.5 |
| Pancreatic ductal adenocarcinoma | Improved | 0.0016 | 15.33 | 22.03 |
| Sarcoma | Improved | 0.033 | 49.27 | 77.47 |
| Cervical squamous cell carcinoma* | Reduced | 0.050 | 39.53 | 28.7 |
| Head-neck squamous cell carcinoma* | Improved | 0.0489 | 48.63 | 58.27 |
| skin cutaneous melanoma (SKCM)@ | Improved | 5.4e-05 | - | - |

The table displays the correlations between *JUNI* expression levels and patient survival in different types of cancer, based on data obtained from Kaplan-Meier Plotter-mediated processing of Pan-cancer RNAseq data. The asterisk (\*) denotes results that were obtained from white populations only. @=
results obtained from GEPIA2 (Tang *et al*, 2019). The table shows the median survival (M.S.) in months for low and high expressers of *JUNI*.

### Short-term silencing of *JUNI* sensitizes cancer cells to chemotherapeutic drugs, whereas prolonged *JUNI* silencing kills cancer cells regardless of challenge exposure

The induction of *JUNI* by DNA damaging agents (Fig 1) suggested that it may play a role in the cellular response to DNA damage. To examine this possibility, *JUNI* was silenced in HMCB, CHL-1 and MDA-MB-231 cells, and 36-40 h later, the cells were either UV irradiated or treated with the chemotherapeutic drugs etoposide or doxorubicin. Cell survival was determined microscopically and using XTT assays. As depicted in Figure 4A, B and Fig. S2, silencing of *JUNI* in all cell lines sensitized them to UV-induced stress and chemotherapeutic drugs. Apoptotic cells exhibiting blebbing membranes and nuclear fragmentation were clearly observed (Fig 4A). In addition, all treatments induced the levels of cleaved caspase 3 more significantly in *JUNI*-silenced cells (Figure 4B). The exposure to UV radiation and the drugs resulted in 30 to 65% reduction in the survival of *JUNI*-silenced cells relative to cells transfected with control siRNA (Fig. 4C and S2). To determine the effects of *JUNI* silencing on sensitivity of cancer cells to chemotherapeutical drugs in another model, spheroids of CHL1 and HMCB cells were generated, exposed to doxorubicin and analyzed for survival 5-8 days later. As depicted in figure 4 D and E the survival of *JUNI*-silenced cells was reduced by 50%, thus supporting *JUNI’s* protective role in cells exposed to DNA damage.

**Figure 4.**
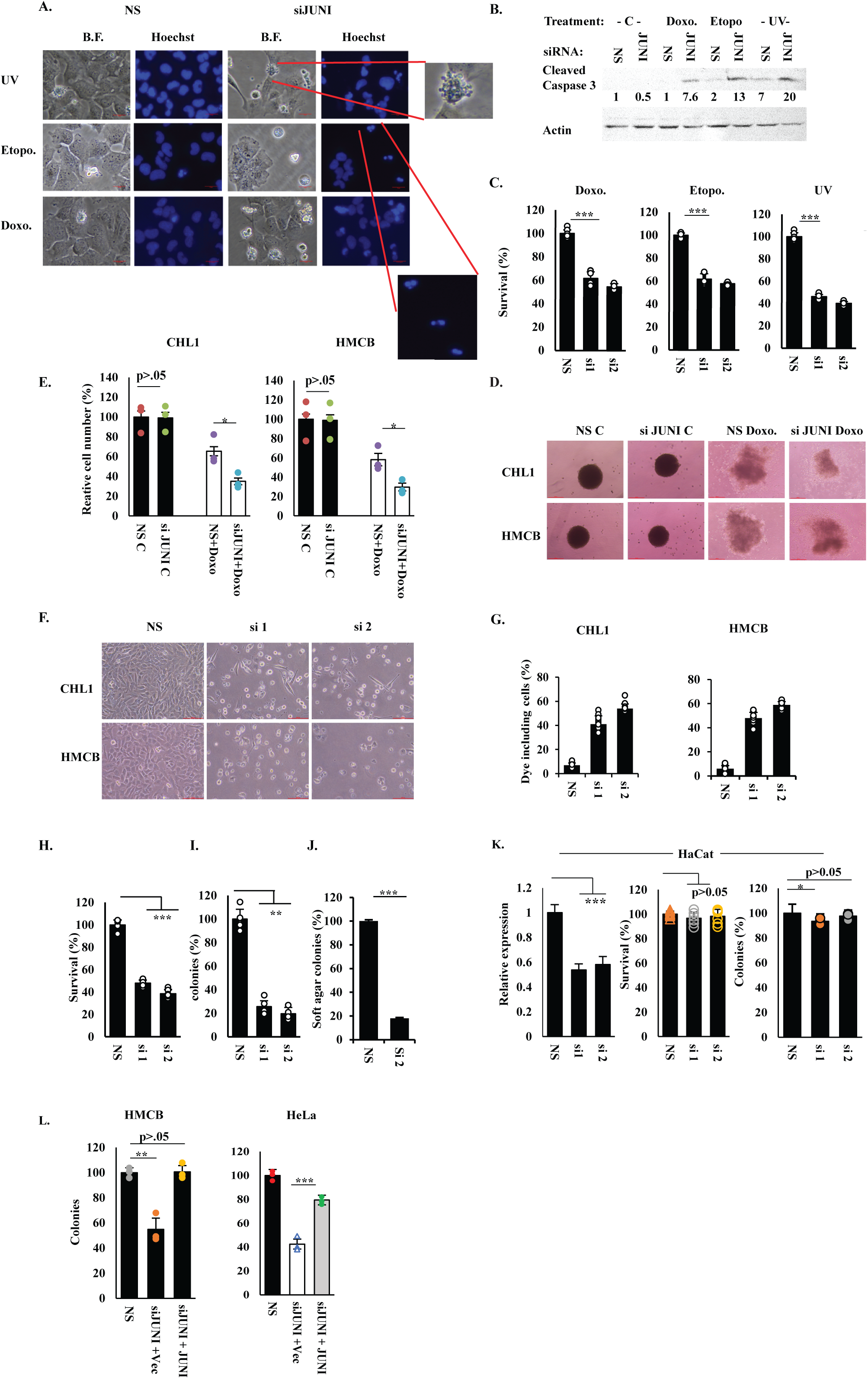
*JUNI* is essential for the survival of cancer cells. A. HMCB cells were transfected with the indicated siRNAs. Thirty-six hours post transfection, the cells were treated with either 20J/m^2^ UVC, 5 μM doxorubicin or 10 μM etoposide and photographed 5- 12 h later. Characteristic morphologies of apoptotic cells are blown up. B. HMCB cells were transfected with the indicated siRNA and treated 36h later with 20J/m^2^ UVC 5 μM doxorubicin and 25 μM etoposide and harvested 12-18 h later to determine the presence of cleaved caspase 3 using specific antibody. Actin was used as a loading control and relative levels are indicated. C. HMCB cells were transfected with the indicated siRNAs and treated 36 h later with 15 J/m^2^ UV, 1μM doxorubicin or 5μM etoposide. XTT was measured 20 h later. Survival of NS -transfected cells was considered 100% in this and in the following XTT experiments. N in all XTT experiments is >3. D-E. Spheroids of the indicated cell lines were exposed to 1-2μM doxorubicin, photographed 5-8 days later (D), collected, tripsinized and counted (E). F. CHL1 and HMCB cells were transfected with the indicated siRNAs and photographed 120 h later. G. Trypan blue-absorbing cells 96 h post transfection with the indicated siRNA were counted and presented as percent of the total population. H. Percentage of cellular survival of HMCB cells measured 120 h post transfection with different siRNAs using XTT. I. Colonies of HMCB cells grown 14 days after transient transfection with NS, si1 or si2. No selection was used. The number of colonies formed by NS-transfected cells was considered 100%. J. HMCB cells were transfected with NS or si2, and 24 h later, the cells were plated into soft agar. Clones in 10 fields were counted 2 weeks later. The number of colonies developed in NS-transfected cells was considered 100%. K. *JUNI* was silenced or not in HaCat cells with the indicated siRNAs. Its expression was determined 36 h post transfection (left panel) using RTqPCR, survival of the cells was determined 120 h post transfection using XTT (middle panel) and 14 days later using colony formation assay (right panel). L. *JUNI* was silenced in the indicated cell line. 24h post siRNA transfection expression vector for the the first exon of *JUNI* was transfected (50-100ng/well in 12 well dish). Transfection was repeated 96 h later and colonies were counted 14 days post plating.

More striking was the fact that *JUNI* silencing for a prolonged period of 96-120 h resulted in elevated cell death even without exposure to additional stress, in all four examined cell lines (Fig. 4 F-H and S3). Unlike the cells exposed to DNA damaging agents the spontaneously dying cells did not exhibit clear apoptotic characteristics and entry of dye into the cells, caused by membrane rupturing, was measured (Fig 4G). The exact causes and mechanisms of cell death are currently under investigation (see discussion), nevertheless, quantification of cell survival using XTT at 120 h post *JUNI* silencing revealed 50-70% cell death in the different cell lines (Fig 4H and S3). The survival rate was further reduced to 10-25% two weeks after silencing, as indicated by the number of colonies that developed in *JUNI*-silenced cells (Fig 4I and S3). Consistent with these results, the silencing of *JUNI* in HMCB and HeLa cells resulted in reduced formation of colonies in soft agar (Fig 4J and S3). The requirement of *JUNI* for survival under these conditions seemed to be specific for cancer cells as its silencing in the non-tumorigenic, spontaneously transformed human skin keratinocyte cell line, HaCaT, did not result in significant cell death (Fig. 4K). Reintroduction of a *JUNI* into *JUNI*-silenced HMCB and HeLa cells rescued them from cell death as reflected by the ability to form colonies in clonogenic assays (Fig. 4L). Overall, we suggest that JUNI is essential for the survival of stress -exposed but also in cancer cells not exposed to DNA damaging agents.

### *JUNI’s* requirement for cellular survival is partially dependent on c-Jun

As c-Jun is a major cellular signaling molecule involved in many aspects of cellular wellbeing, cell death after *JUNI* silencing can potentially be caused indirectly by the decline in c-Jun levels. To address this possibility, we compared the effects inflicted on survival after *JUNI* silencing to those observed after *JUN* silencing. We used specific siRNAs to silence either *JUNI* or *JUN* in 3 different cell lines, MDA-MB-231, HeLa and HMCB (Figure 5). Interestingly, although *JUN* mRNA levels in cells transfected with specific *JUN* siRNA were at least 20% lower than those in *JUNI* silenced cells (Fig 5 A-C, Expression), their survival 96 h after *JUN* silencing was five to 30% higher (Fig 5 A-C, Survival XTT/short). This difference was extenuated two weeks later. A reduction of 45-60% in colonies formed after *JUNI* silencing relative to *JUN* silenced cells was observed (Fig 5 A-C, Survival colonies/long). These results demonstrate that in the examined cell lines, *JUNI* is more important for cellular survival than *JUN*. As demonstrated before (Fig. 2), *JUN* silencing had no significant effect on *JUNI* expression (Fig. 5 A-C, Expression).

**Figure 5.**
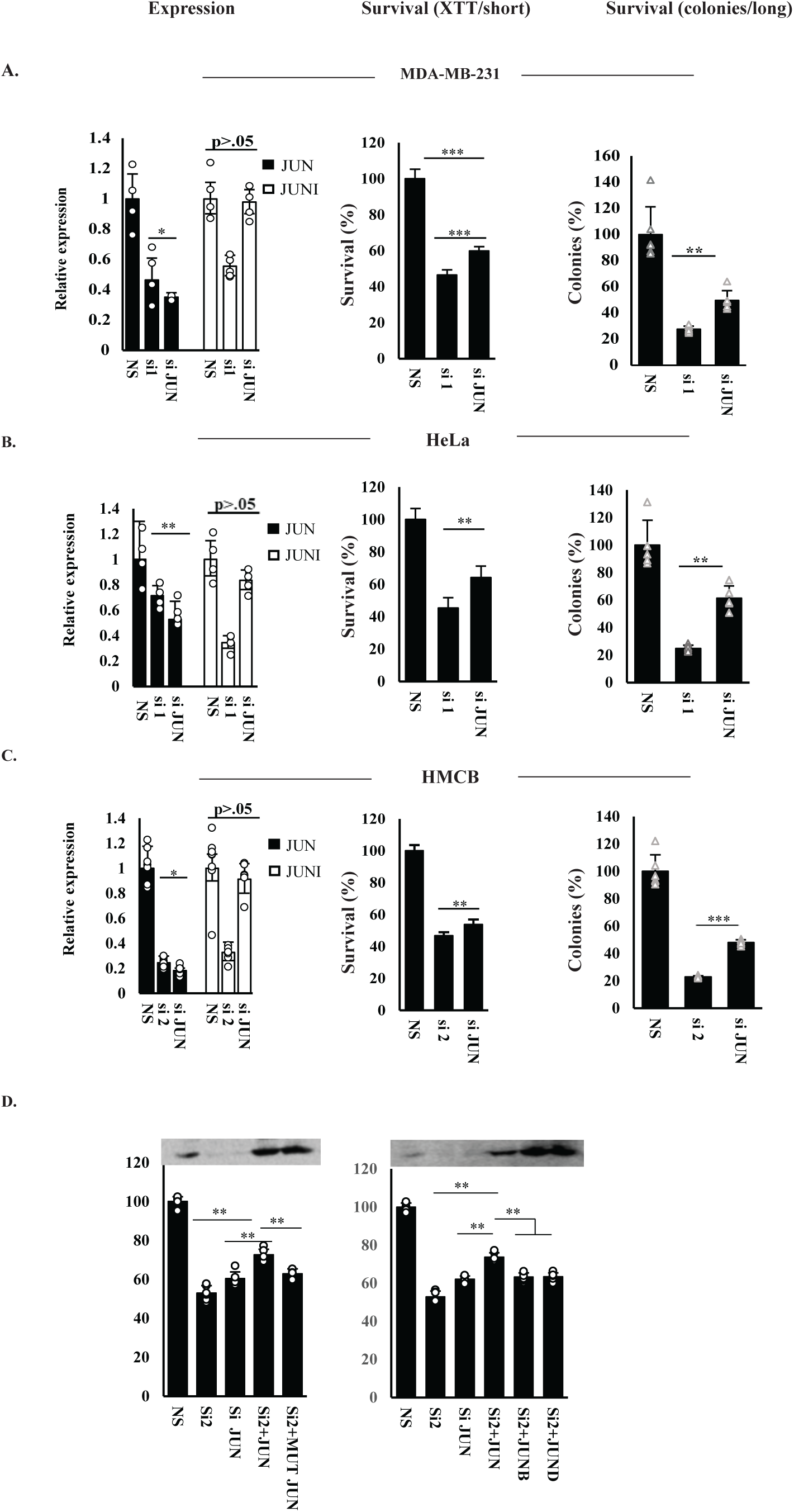
*JUN* mediates only part of *JUNI’s* effects on survival. MDA-MB-231 (A) HeLa (B) and HMCB cells (C) were transfected with NS, siRNA against *JUNI* or siRNA against *JUN*. The levels of *JUN* (black bars) and *JUNI* (white bars) were determined 30 h after transfection using RT-qPCR (left panels). Short-term effects on the viability of the transfected cells were determined using XTT 96 h later in HeLa cells or 120 h later in the other cell lines (Middle panels, Survival XTT/short). Long-term effects on viability were measured two weeks post transfection using a clonogenic assay (Right panel, survival colonies/long). D. HMCB cells silenced for *JUNI* were cotransfected with the indicated expression vectors. Cell survival was examined using the XTT assay 96 h post transfection. c-Jun protein expression levels in the transfected cells are presented.

To further test the contribution of c-Jun to *JUNI*’s pro-survival activity, we performed rescue experiments in which *JUNI* was silenced in HMCB cells and exogenous c-Jun was expressed to complement the repressed endogenous gene. *JUNI* silencing reduced cell survival by 50%, whereas *JUN* silencing reduced cell survival by 40% (Fig. 5D). Restoration of *JUN* expression improved the survival of *JUNI*-silenced cells. However, the rescue was partial (Fig. 5D). Mutant c-Jun that cannot bind DNA (272/273E) (Yogev *et al*., 2008) or other *JUN* family members, *JUNB* or *JUND*, could not rescue cell death caused by *JUNI* silencing, thus, suggesting a specificity for transcriptional active c-Jun in the rescue. These results indicate that *JUNI’*s effects on cellular survival are partially dependent on c-Jun expression and predict that additional *JUNI*-regulated factors are essential for cellular survival.

### Antagonistic interactions of *JUNI* with DUSP14 mediate c-Jun induction by UV and survival

To explore the identity of protein interactors that may affect the ability of *JUNI* to regulate c-Jun expression and affect cell death, we identified interacting protein partners of *JUNI* by applying the incPRINT screen (Graindorge *et al*., 2019). Using the shortest yet stable isoform of *JUNI* that contained the *JUN*-affecting sequences as RNA bait, we identified 57 *JUNI*-interacting proteins (Fig. 6A). Analyses of their cellular functions revealed enrichment of proteins involved in various fundamental, basic aspects of cellular wellbeing, such as splicing, ribosome biosynthesis, and mitosis (Table S2). Interestingly, c-Jun itself does not interact with JUNI (Table S2, Normalized luciferase intensity MS2, RLU =0.44). By contrast, the dual specificity protein phosphatase 14 (DUSP14; also known as MKP6) was one of the highly rated interacting proteins with *JUNI* (Fig 6A, B). DUSP14 can directly dephosphorylate or indirectly limit the phosphorylation of MAPKs essential for c-Jun expression: JNK, p38 and ERK (Marti *et al*, 2001; Yang *et al*, 2014). We validated the interaction between DUSP14 and *JUNI* in HeLa cells by crosslinking and immunoprecipitation (CLIP). We examined the enrichment of endogenous *JUNI* following immunoprecipitation with transfected DUSP14 or GFP. Specificity was determined by comparison of *JUNI* enrichment to other unrelated nuclear lncRNAs *MALAT1* and *PVT1*. These experiments demonstrated a specific association of *JUNI* with DUSP14 (Fig. 6C). As *JUNI* is required for efficient c-Jun upregulation after UV exposure and DUSP14 inhibits JNK, a stress-activated, positive regulator of *JUN* transcription and protein expression (Davis, 2000; Karin & Gallagher, 2005), we hypothesized that *JUNI* antagonizes DUSP14 activity to enable full-scale c-Jun induction post-stress exposure and that the antagonism may have a role in other *JUNI*-dependent activities discovered in this study. The potential *JUNI*-DUSP14 antagonism was examined at 3 levels. Biochemically, we sought to test if *JUNI* enables efficient JNK and c-Jun phosphorylation, thus resulting in c-Jun induction post UV exposure. At the cellular level, we examined whether *JUNI*- DUSP14 antagonism is reflected by effects on cellular survival, and most importantly, we examined whether the expression of both inversely correlates with the survival of human patients suffering from cancer types in which *JUNI* levels correlate with survival (Table 1).

**Figure 6.**
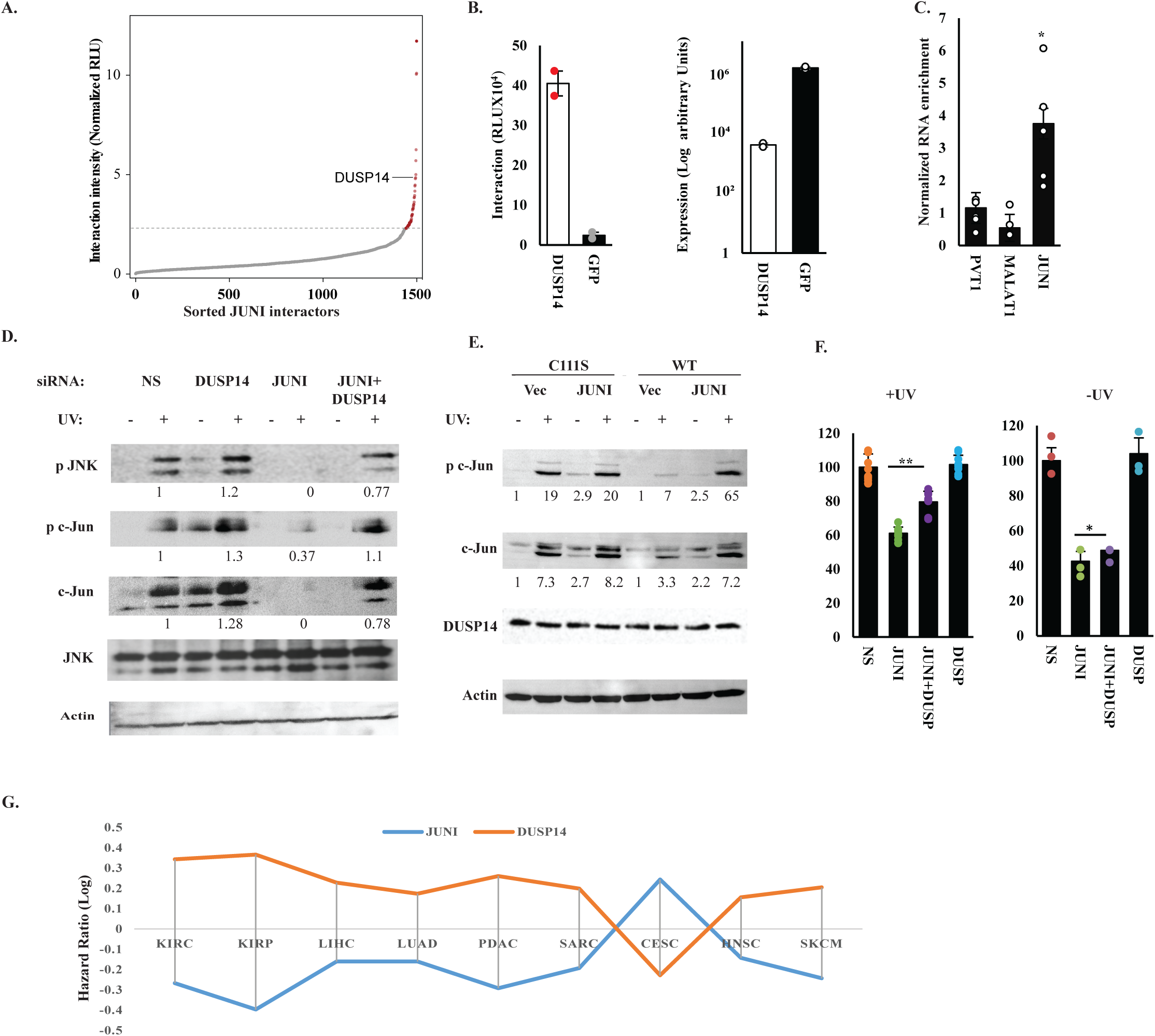
*JUNI* interacts with and antagonizes DUSP14 activity biochemically and in cancer. A. Normalized *JUNI*-protein interaction intensities averaged from two incPRINT biological replicates, sorted in increasing order. The horizontal dotted line represents the interaction intensity cutoff used for the classification of *JUNI* interactors. Red dots are *JUNI*-interacting proteins; gray dots are proteins that do not bind to *JUNI*. See the ‘Methods’ section for data normalization. RLU are relative light units. B. Intensities of *JUNI* interactions with DUSP14 and GFP were obtained from two biological replicates of the incPRINT screen. On the left, interaction intensities detected between *JUNI* and the indicated proteins. On the right, the expression level of each protein was measured using ELISA. C. HeLa cells were transfected with DUSP14 or GFP. UV crosslinking immunoprecipitation was performed. The ratio of RNAs coprecipitated with DUSP14 relative to RNAs in the whole cell extract was further normalized to the ratio obtained after GFP precipitation. Enrichment of the indicated lncRNAs is depicted. D. HMCB cells were transfected with the indicated siRNAs, irradiated with 30 J/m 48 h later and harvested 5 h post irradiation. Levels of the indicated proteins were measured by immunoblotting using specific antibodies as described before. E. HMCB cells were transfected with the indicated plasmids and 24 h later irradiated or not with 25 J/m^2^ UV. Cells were harvested 4 h later and the levels of the indicated proteins were measured using specific antibodies. DUSP14 was measured using HA antibody. F. HMCB cells were transfected with the indicated siRNAs, UV irradiated 40 h later or not. Irradiated cells were harvested 12 h after irradiation, whereas non-irradiated cells were harvested 96 h post transfection. Viability was determined using XTT. G. Comparison of hazard ratios of *JUNI* and DUSP14 in the indicated human cancers. Data obtained from Kaplan-Meier Plotter-mediated processing of Pan-cancer RNAseq data (Nagy *et al*., 2021).

To explore the effects on c-Jun, we silenced DUSP14, *JUNI* or both in HMCB cells that were later exposed to UV radiation and JNK activation, c-Jun phosphorylation and its total level were determined. Indeed, silencing *JUNI* inhibited JNK phosphorylation (pThr183/pTyr185) in UV-treated cells to a nearly undetectable level, and consequently, Ser63 phosphorylation of c-Jun therefore, prevented its induction by UV (Fig. 6D). Co-silencing of DUSP14 together with *JUNI* restored JNK and c-Jun phosphorylation to levels silhgtly lower than those observed in non-silenced cells. To prove that the effect of *JUNI* on c-Jun induction is mediated by DUSP14 phosphatase activity we transfected HMCB cells with wt or phosphatase dead (C111S) DUSP14 together with *JUNI* expression vector or an empty one. Cells were irradiated 24h post transfection and the phosphorylation as well total levels of c-Jun were monitored (Fig 6E). As depicted in Fig 6E, *JUNI* considerably elevated c-Jun phosphorylation and expression only in cells transfected with phosphatase active DUSP14. These data suggest that *JUNI* regulates JNK phosphorylation and c-Jun levels by antagonizing DUSP14 activity.

To assess the importance of DUSP14 in *JUNI*-dependent effects on the survival of stress-exposed cells, *JUNI*, DUSP14, or both were silenced in HMCB (Fig 6F) and HeLa cells (Fig S4) that were later exposed to UV, and cell survival was measured. Consistent with the previously indicated results (Fig 4), *JUNI*-silenced cells were 40% more sensitive to radiation than non-silenced cells (Fig. 6F and S4). DUSP14 silencing did not significantly improve the survival of irradiated cells expressing normal levels of *JUNI*. Remarkably, it improved the survival of *JUNI*-silenced cells by 29 and 33% in HMCB and HeLa cells, respectively (Fig. 6F and S4), suggesting that *JUNI*’S expression is important for antagonizing DUSP14 in order to improve the survival of stress-exposed cells. Nevertheless, the negative effect of *JUNI* silencing on the survival of stressed cells was not fully rescued by repression of DUSP14, suggesting the presence of additional targets. In contrast, the rescue from cell death caused by prolonged silencing of *JUNI* in non-irradiated cells was considerably lower, suggesting that the impact of antagonistic interactions between *JUNI* and DUSP14 is most significant in stress-exposed cells (Fig. 6F and S4).

### Expression levels of *JUNI*-interacting proteins in tumors predict prognosis value in clear cell renal cell carcinoma (ccRCC)

To test the hypothesis that *JUNI*-DUSP14 antagonism has a potential role in human cancer, we analyzed the hazard ratios (HR) of *JUNI* and DUSP14 expression levels in the 11 types of cancer in which *JUNI* expression correlated significantly with disease prognosis (Table 1). Interestingly, a strict inverse relationship was observed in nine out of the eleven (Fig. 6G). The inverse correlation was reflected not only in the effect on survival but also in magnitude of the effect, thus excluding randomness. In line with hazard ratios, the median survival associated with expression levels was also coherently inverse in eight out of these 11 cancers (Table S4). The association of *JUNI* with improved survival and DUSP14 with reduced one was particularly significant in ccRCC (p=5.7E-05 and p=2.9E-05 for *JUNI* and DUSP14 respectively) (Fig 7A). In this cancer, high levels of *JUNI* were associated with 87% prolonged median survival, whereas high levels of DUSP14 were associated with a 55% reduction in the median survival of these patients (Table S3). Analysis of *JUNI* expression in ccRCC tumors revealed that in accordance with an assumed negative role in ccRCC tumorigenicity, it’s levels were significantly reduced in the tumors relative to the normal tissues (p=1.2E-11), whereas DUSP14 expression was increased in a very significant manner (p=1.0E-20) (Fig 7B). These results suggest that *JUNI* has a possible role in ccRCC pathogenesis and that, similar to cell lines, it antagonizes DUSP14 also in this cancer type.

**Figure 7.**
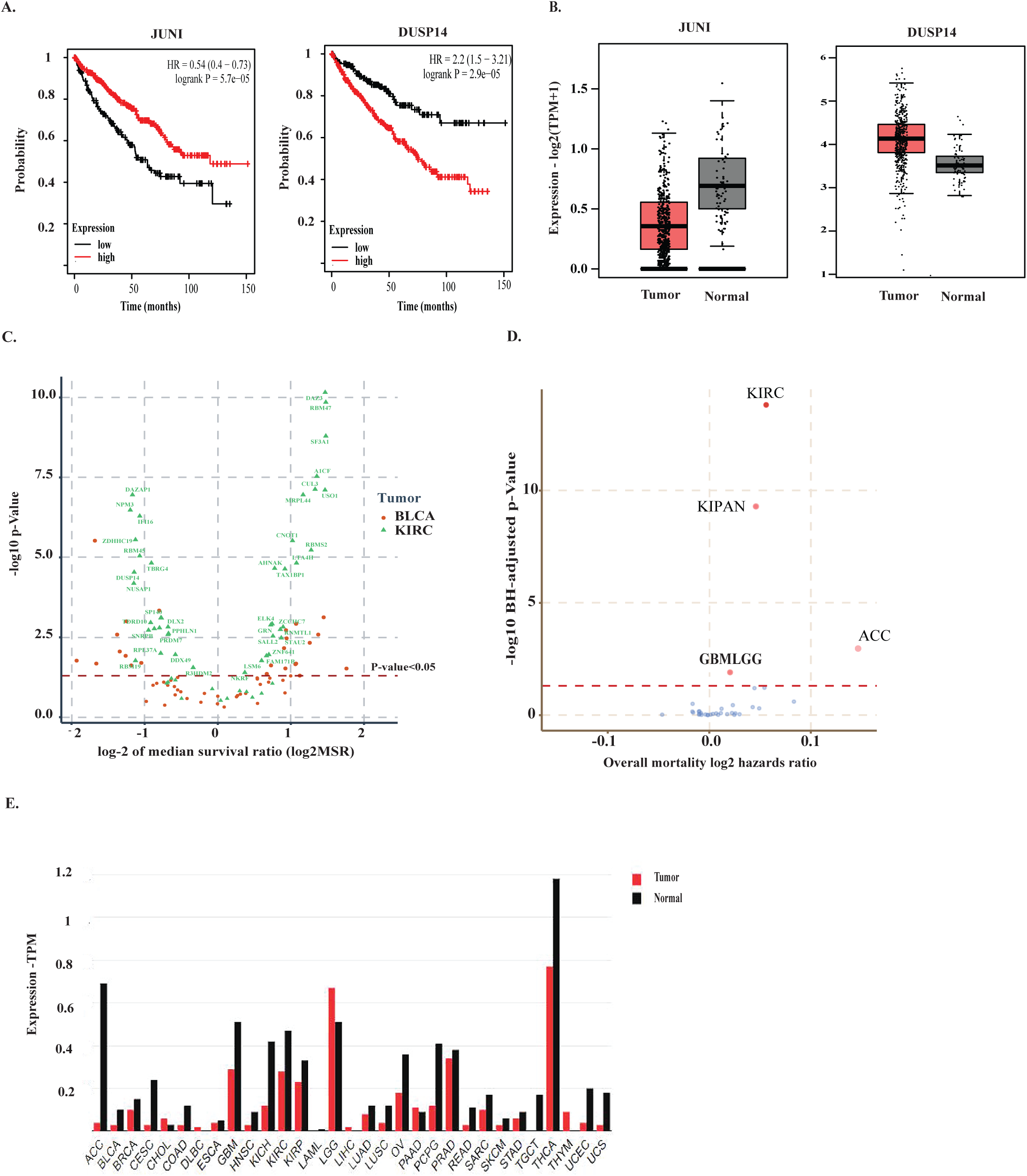
Levels of *JUNI* and its interacting proteins are prognostic factors in ccRCC. A. Kaplan-Meier plots of patients expressing high (red) or low (black) levels of the indicated RNAs in ccRCC tumors (Data source identical to 6G). B. *JUNI* and DUSP14 expression in ccRCC samples and normal tissues was analyzed using TCGA and GTX data and GEPIA tool (Tang *et al*., 2017). C. Volcano plot showing the median survival ratios (MSR) and -log10 p-values of the 55 incPRINT-derived genes in patients with ccRCC (KIRC) or bladder cancer (BLCA). The MSR values are shown in the x-axis and the -log10 of the log rank p-values are shown in the y-axis. Data were extracted from the Pan-cancer RNA-seq Kaplan-Meier plotter (Nagy *et al*., 2021). D. A volcano plot displaying the risk of mortality associated with a change in the expression of the 55 incPRINT-derived genes across various TCGA datasets. The x-axis shows the degree of this risk (measured as log2 hazard ratios), while the y-axis shows the statistical significance of these findings (measured as -log10 of adjusted p-values). These analyses cover different cancer types, including clear cell renal cell carcinoma (KIRC), pan-kidney cohort (KIPAN), adrenocortical carcinoma (ACC), glioblastoma multiforme and low-grade glioma (GBMLGG). E. Analysis of *JUNI* levels in normal and tumor samples. Data obtained from the GEPIA (Tang *et al*., 2017), which uses a TCGA and GTEx RNA sequencing expression data.

Next, we examined whether the prognostic significance in ccRCC is unique to DUSP14 or shared with other *JUNI*-interacting proteins. To that end we determined the prognostic significance of each of the top 55 *JUNI*-interacting proteins by analyzing Pan-cancer RNAseq data for ccRCC with Kaplan-Meier plotter and comparing these values to the prognostic significance of the same protein array in bladder cancer (BLCA), another urinary-system cancer (Figure 7C). Two important observations were found; the portion of proteins which are significantly linked with prognosis in ccRCC was about 76% compared with only 43% in bladder carcinoma and, moreover, the significance values were much higher (Fig. 7C), thus suggesting specificity for ccRCC. To further address the specificity of the entire *JUNI*-interacting-proteins-array, identified using incPRINT, with ccRCC prognosis, a combined expression score was driven from all 55 proteins (see Methods), and analyzed for prognostic significance across 34 Cancer Genome Atlas (TCGA) cohorts (cancer types). As depicted in Figure 7D the expression of genes coding for *JUNI*-interacting proteins provides a unique signature with an extremely high significance in the ccRCC cohort followed only by the pan-kidney cohort (KIPAN) and adrenocortical carcinoma (ACC), both kidney-related. Minor but significant association of this gene set with prognosis was also observed in glioma patients (GBMLGG cohort) (Fig 7D). This gene combination was not significant for prognosis in neither of the other cancer types examined. Analysis of *JUNI*’s expression in tumors and normal samples from TCGA and the GTEx data bases using the GEPIA web server(Tang *et al*, 2017) revealed that *JUNI*’s levels are reduced in all kidney and kidney related cancers, most significantly in ACC (Fig 7E).

## Discussion

Overall, we discovered a novel lncRNA that plays fundamental roles in cellular survival. The major concern that every cellular effect observed post-silencing of *JUNI* is derived from the regulation of c-Jun was ruled out, as *JUNI* seems to be more potent than c-Jun in maintaining the survival of the cancer cells examined (Fig. 5). Furthermore, exogenous c-Jun only partially rescued HMCB cells silenced for *JUNI*, demonstrating that cell death is partially independent of c-Jun. Therefore, we suggest that much of the biological effects attributed to *JUN*I are c-Jun independent and presumably occur by regulating the activity of interacting proteins such as DUSP14.

The nonspecific inhibition of transcription initiation (Rockx *et al*, 2000) and elongation (Williamson *et al*, 2017) observed after UV exposure suggests that *JUNI* induction under stress is specific. The strongest effects of *JUNI* on c-Jun regulation were observed after stress exposure. *JUNI* appears to respond in a very rapid manner to UV exposure, as required from an inducer of c-Jun, an “immediate early protein”. Its induction requires JNK and p38 activation, and in turn, it binds and antagonizes DUSP14, a negative regulator of these kinases, in a positive feedback loop. Although regulating the major JNK-c-Jun cellular pathway post DNA damage is supported by the ability of *JUNI* to induce c-Jun expression *in trans* and to enable efficient JNK phosphorylation after UV exposure, it does not necessarily exclude additional effects, *in cis,* of *JUNI* on c-Jun expression in untreated cells via association with other interacting proteins. Both the effects of *JUNI* on c-Jun induction and cellular survival were demonstrated using underexpression conditions by targeting the common, first exon, of *JUNI.* Moreover, this exon was also sufficient for c-Jun induction upon stress exposure, under conditions of overexpression.

*JUNI* is essential for cell survival in both chemotherapy-treated and untreated cells. We predict that the causes and mechanisms of death of cells exposed to stress are different from those in cells that are not. Sensitization of all cell types examined in this study to chemotherapeutic drugs occurs days prior to spontaneous cell death. The morphology of the spontaneous cell death is not apoptotic, and the *JUNI*- interacting protein DUSP14 inhibits more efficiently stress-dependent cell death in comparison to cell death observed after prolonged silencing of *JUNI* (Figure 6F and S4). We predict that the cause of spontaneous cell death following *JUNI* silencing is not singular. Examination of the plethora of functions of the putatively interacting proteins may suggest that prolonged silencing leads to abnormalities in different cellular functions culminating in synthetic cell death.

As JNK activity usually elevates cell death post UV exposure (Tournier *et al*, 2000), the ability of DUSP14 silencing to rescue it upon JUNI silencing is surprising. Three potential mechanisms may account for the rescue. First, DUSP14 inhibition increases the activity of protective proteins that are negatively regulated by it. For example, NFκB is downstream of TAK1-TAB1 proteins which are negatively regulated by DUSP14(Yang *et al*., 2014). Second, low constitutive JNK activation may induce autophagy which exerts protective activity (Sample & He, 2017; Wu *et al*, 2019). Finally, the indirect suppression of genes involved in EMT may lead to a less mesenchymal phenotype reflected by reduced motility. Given that EMT is associated with increased drug resistance (Shibue & Weinberg, 2017), it is possible that the mesenchymal to epithelial transition (MET) process occurring in these cells after silencing *JUNI* sensitizes the cells to chemotherapeutic drugs post *JUNI* silencing.

We also examined the correlation between *JUNI* expression and the prognosis of ccRCC. Our results demonstrate that high levels of *JUNI* are associated with longer survival in ccRCC patients (Fig 7 and Table 1). Additionally, the high prognostic significance levels of 76% of its newly discovered interacting proteins suggest that *JUNI*’s activity may be mediated via the regulation of these proteins (Fig 7). This finding suggests that the combined-expression-analysis of these proteins may provide a prognostic value that is more specific to ccRCC than any other cancer. This is the first observation connecting *JUNI* and its interacting genes to ccRCC. Its potential role in this disease opens a novel direction for the involvement of this lincRNA in ccRCC.

## Acknowledgments and funding sources

## Acknowledgements

E.S. and V.K. are funded by the Israel Cancer Association, 20220102. A.S is supported by grants from Institut Curie, LabEx DEEP (ANR-11-LABX-0044, ANR-10-190 IDEX-0001-02) and X.S.C. is supported by a Foundation pour la Recherche Médicale doctoral fellowship.

F.K.M. and M.D. were supported by a UK Medical Research Council Career Development Award (MR/P009417/1) to F.K.M.

